# FGF21 has a sex-specific role in calorie-restriction-induced beiging of white adipose tissue in mice

**DOI:** 10.1101/2022.07.29.501882

**Authors:** Mariah F. Calubag, Ismail Ademi, Chung-Yang Yeh, Reji Babygirija, Alyssa M. Bhoopat, Ildiko Kasza, Michelle M. Sonsalla, Cara L. Green, Heidi H. Pak, Dudley W. Lamming

## Abstract

Calorie restriction (CR) promotes healthspan and extends the lifespan of diverse organisms, including mice, and there is intense interest in understanding the molecular mechanisms by which CR functions. Some studies have demonstrated that CR induces fibroblast growth factor 21 (FGF21), a hormone that regulates energy balance and that when overexpressed, promotes metabolic health and longevity in mice, but the role of FGF21 in the response to CR has not been fully investigated. We directly examined the role of FGF21 in the physiological and metabolic response to a CR diet by feeding *Fgf21*^*-/-*^and wild-type control mice either *ad libitum* (AL) diet or a 30% CR diet for 15 weeks. Here, we find that FGF21 is largely dispensable for CR-induced improvements in body composition and energy balance, but that lack of *Fgf21* blunts CR-induced changes aspects of glucose regulation and insulin sensitivity in females. Surprisingly, despite not affecting CR-induced changes in energy expenditure, loss of *Fgf21* significantly blunts CR-induced beiging of white adipose tissue in male but not female mice. Our results shed new light on the molecular mechanisms involved in the beneficial effects of a CR diet, clarify that FGF21 is largely dispensable for the metabolic effects of a CR diet, and highlight a sex-dependent role for FGF21 in the molecular adaptation of white adipose tissue to CR.

## Introduction

Calorie restriction (CR), defined as a dietary regimen in which calories are reduced without malnutrition, promotes healthspan and increases lifespan in diverse organisms ranging from yeast to mice and non-human primates (Green, Lamming, & Fontana, 2022; Lin, Defossez, & Guarente, 2000; Mattison et al., 2017; McCay, Crowell, & Maynard, 1935). CR promotes metabolic health in mammals, reducing adiposity and improving blood sugar control (Mattison et al., 2017; Pak et al., 2021; Wei et al., 2019; Yu et al., 2019). Despite almost a century of effort, the physiological and molecular mechanisms by which CR functions are still not totally understood, stymieing efforts to develop CR mimetics which could promote health in the rapidly aging global populace.

As CR works to slow aging in all tissues, it has been suggested that CR may work in part through endocrine factors. The role of hormones, including growth hormone, insulin, insulin-like growth factor 1 (IGF-1) and adiponectin, have been explored by numerous groups, but evidence that one of these hormones is responsible for the beneficial effects of CR remains elusive (Balasubramanian et al., 2021; Bonkowski, Rocha, Masternak, Al Regaiey, & Bartke, 2006; Yu et al., 2019). One endocrine factor that could play a role in CR that has not yet been fully explored is fibroblast growth factor 21 (FGF21).

FGF21 is a hormone produced by multiple tissues including the liver and white adipose tissue (WAT) in response to a diverse array of nutrient stresses, including fasting, protein restriction, and restriction of specific dietary amino acids (Laeger et al., 2014; Nishimura, Nakatake, Konishi, & Itoh, 2000; Yap et al., 2020; Yu et al., 2021). FGF21 promotes insulin sensitivity and regulates energy balance by promoting food consumption, activating brown adipose tissue (BAT) and stimulating the beiging of inguinal white adipose tissue (iWAT). FGF21 mediates the adaptive starvation response to induce ketogenesis, gluconeogenesis, lipolysis, and lipid β-oxidation (Coskun et al., 2008; Inagaki et al., 2007; Izumiya et al., 2008; Xu et al., 2009). In addition to mimicking many of the beneficial metabolic effects of a CR diet, overexpression of FGF21 significantly extends lifespan (Zhang et al., 2012). Intriguingly, several studies have found that FGF21 is induced by CR in mice and rats (Fujii et al., 2019; Thompson et al., 2014), though to different extents; we also have observed an effect of CR on blood levels of FGF21.

Here, we directly examine the role of FGF21 in the metabolic response to CR by examining the effects of CR in wild-type and *Fgf21*^*-/-*^ mice of both sexes. Surprisingly, we find that circulating levels of FGF21 are not elevated by CR in the fed state, and that FGF21 is largely dispensable for the metabolic benefits of a CR diet in both males and females. Both wild-type and *Fgf21*^*-/-*^ mice placed on CR become lean, glucose tolerant, and insulin sensitive, with a minor effect of *Fgf21* deletion on fasting blood glucose levels in female mice. Similarly, we find that FGF21 is not required for the effects of CR on energy balance. Surprisingly, we find that loss of *Fgf21* blunts the CR-induced reprogramming of white adipose tissue metabolism in male mice. We conclude that despite the similarity in FGF21-induced and CR-induced phenotypes, and our discovery of a role for FGF21 in the CR-induced reprogramming of white adipose tissue, FGF21 is largely dispensable for the effects of CR on metabolic health.

## Materials and Methods

### Animal care, housing and diet

All procedures were performed in conformance with institutional guidelines and were approved by the Institutional Animal Care and Use Committee of the William S. Middleton Memorial Veterans Hospital. Male and female wild-type and *Fgf21*^*-/-*^ mice were generated by crossing CMV-Cre mice (Schwenk, Baron, & Rajewsky, 1995) from the Jackson Laboratory (006054) with mice expressing a floxed allele of *Fgf21* (Potthoff et al., 2009) from The Jackson Laboratory (022361), and then crossed with C57BL/6J mice to remove CMV-Cre. All mice were acclimated to the animal research facility for at least one week before entering studies. All animals were housed in static microisolator cages in a specific pathogen-free mouse facility with a 12:12 h light–dark cycle, maintained at approximately 22 °C.

At approximately 9 weeks of age, all animals were singly housed and placed on 2018 Teklad Global 18% Protein Rodent Diet for 1 week before randomization. At 10 weeks of age, mice were randomized to either an *ad libitum* (AL) diet or 30% calorie restricted (CR) diet, in which animals were fed once per day during the beginning of the light period, at approximately 7am. A stepwise reduction in food intake starting at 20% was carried out for mice in the CR group in week 1 before maintenance of a 30% restriction. The caloric intake of the mice in the AL group was calculated weekly to determine the appropriate number of calories to feed the mice in the CR groups. This feeding schedule has been widely used and extends the lifespan of mice (Pak et al., 2021; Turturro et al., 1999; Yu et al., 2019).

### Metabolic Phenotyping

Glucose, insulin and pyruvate tolerance tests were performed by fasting all mice for 7 hours or overnight (∼21-22 hours) and then injecting either glucose (1g/kg), insulin (0.5U/kg) or pyruvate (2g/kg) intraperitoneally (Bellantuono et al., 2020; Yu et al., 2019). Glucose measurements were taken using a Bayer Contour blood glucose meter and test strips. Mouse body composition was determined using an EchoMRI Body Composition Analyzer. For assay of multiple metabolic parameters (O_2_, CO_2_, food consumption, and activity tracking), mice were acclimatized to housing in a Columbus Instruments Oxymax/CLAMS-HC metabolic chamber system for approximately 24 hours, and data from a continuous 24-hour period was then recorded and analyzed. AL-fed animals had *ad libitum* access to diet; CR groups were fed once per day at the beginning of the light cycle.

### Collection of tissues for molecular and histological analysis

For analysis of plasma FGF21 levels in Figure 1A, mice were sacrificed at 7AM prior to the regularly scheduled morning feeding time of the CR-fed animals. For all other experiments, mice were euthanized in the fed state after approximately 15 weeks. Mice euthanized in the fed state had all food removed starting at 3pm the day prior to sacrifice, fed at 7am the day of sacrifice, and then euthanized 3 hours later. Following blood collection via submandibular bleeding, mice were euthanized by cervical dislocation and most tissues were rapidly collected, weighed and then snap frozen in liquid nitrogen. A portion of iWAT was fixed in 10% formalin for 48 hours before being transferred to 70% ethanol, sectioned, and Hematoxylin and eosin stained by the UWCCC Experimental Pathology Laboratory. Images of the iWAT were taken using an EVOS microscope (Thermo Fisher Scientific Inc., Waltham, MA, USA) at a magnification of 40X as previously described (Cummings et al., 2018; Yu et al., 2021). Scale bars were inserted automatically or manually by the investigator. For quantification of adipocyte size, six independent fields were obtained for each tissue from each mouse and quantified using ImageJ (NIH, Bethesda, MD, USA). For quantification of multilocular cells, a blinded participant counted the number of multilocular cells per image for each mouse using ImageJ (NIH, Bethesda, MD, USA).

**Figure 1.**
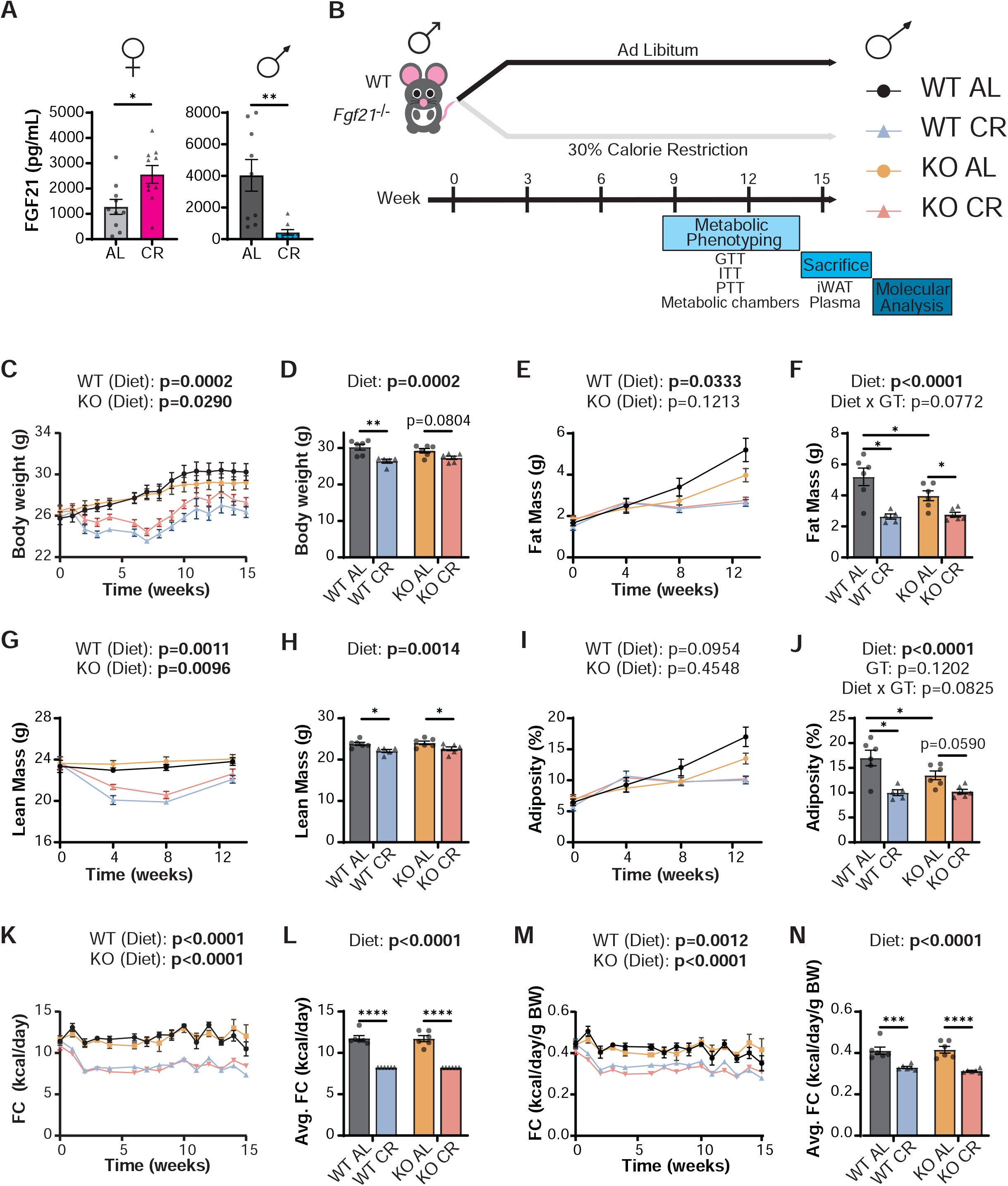
Loss of *Fgf21* does not impact the effect of CR on body weight, body composition and food consumption in male mice. (A) Plasma FGF21 levels from female and male mice on either an AL or CR diet for 20 weeks (B) Experimental design. (C-D) Body weight measurement of male mice on the indicated diets (B) with final body weight at 15 weeks (C). (E-F) Fat mass measurement of male mice on the indicated diets (E) with final fat mass at 13 weeks (F). (G-H) Lean mass measurement of male mice on the indicated diets (G) with final lean mass at 13 weeks (H). (I-J) Adiposity of male mice on the indicated diets (I) with final adiposity at 13 weeks (J). (K-L) Food consumption (kcal/day) of male mice on the indicated diets (K) with average food consumed (L). (M-N) Food consumption per gram of body weight (kcal/day/g BW) of male mice on the indicated diets (M) with average food consumed per gram of body weight (N). (A) n=10 females/group, n=9 males/group, *p=0.0122, **= p=0.0027, unpaired two-tailed t-test. (C-N) n=5-6 mice/group. For longitudinal studies, statistics for the overall effects of genotype, diet, and the interaction represent the p value from a two-way repeated measures (RM) ANOVA or residual maximum likelihood (REML) analysis conducted individually for each genotype. For analyses of weight or body composition as a single time point, or analysis of average food consumption, statistics for the overall effects of genotype, diet, and the interaction represent the p value from a two-way ANOVA; *p<0.05, **p<0.01, **p<0.001, ****p<0.0001 Sidak’s test post 2-way ANOVA. Data represented as mean ± SEM.

### Quantitative real-time PCR

RNA was extracted from liver or iWAT using TRI Reagent according to the manufacturer’s protocol (Sigma-Aldrich). The concentration and purity of RNA were determined by absorbance at 260/280 nm using Nanodrop (Thermo Fisher Scientific). 1 μg of RNA was used to generate cDNA (Superscript III; Invitrogen, Carlsbad, CA, USA). Oligo dT primers and primers for real-time PCR were obtained from Integrated DNA Technologies (IDT, Coralville, IA, USA); sequences are in Table S1. Reactions were run on an StepOne Plus machine (Applied Biosystems, Foster City, CA, USA) with Sybr Green PCR Master Mix (Invitrogen). Actin was used to normalize the results from gene-specific reactions.

### ELISA assays and kits

Blood plasma for insulin was taken the morning of euthanasia from fasted mice prior to refeeding as well as from the terminal submandibular bleeding. Blood plasma for FGF21 was obtained from the terminal submandibular bleeding. Plasma insulin was quantified using an ultra-sensitive mouse insulin ELISA kit (90080), from Crystal Chem (Elk Grove Village, IL, USA). Blood FGF21 levels were assayed by a mouse/rat FGF-21 quantikine ELISA kit (MF2100) from R&D Systems (Minneapolis, MN, USA).

### Statistics

Data are presented as the mean ± SEM unless otherwise specified. Statistical analyses were performed using one-way or two-way ANOVA followed by Tukey–Kramer post hoc test, as specified in the figure legends. Other statistical details are available in the figure legends. In all figures, n represents the number of biologically independent animals. Sample sizes were chosen based on our previously published experimental results with the effects of dietary interventions (Cummings et al., 2018; Fontana et al., 2016; Yu et al., 2021; Yu et al., 2019; Yu et al., 2018). Data distribution was assumed to be normal, but this was not formally tested. Phenotypic data (i.e. body weight, body composition, metabolic health) analyses for PCA plots were performed in R (v. 4.0.3) using gplots. PCA plots were produced by imputing missing data and scaling the data using the R package “missMDA” using the PCA analysis and plots were generated using the R packages “factoextra” and “factoMine” (Husson, Josse, & Lê, 2008; Kassambara & Mundt, 2020). For the heatmap, gene expression data was logged to the base 2 before the plot of generated (Warnes et al., 2005). PCA plots were produced by imputing missing data and scaling the data using the R package “missMDA” using the PCA analysis and plots were generated using the R packages “factoextra” and “factoMine” (Husson et al., 2008; Kassambara & Mundt, 2020).

### Randomization

All studies were performed on animals or on tissues collected from animals. Animals of each sex and strain were randomized into groups of equivalent weight before the beginning of the *in vivo* studies.

## Results

### Loss of FGF21 does not affect the response of body weight and composition to a CR diet

The effect of CR on FGF21, as well as the role of FGF21 in the response to CR, is unclear (Fujii et al., 2019; Thompson et al., 2014). We determined FGF21 levels in the blood of both male and female *ad libitum* (AL)-fed mice and mice in which calories were restricted by 30%. We observed a significant increase of FGF21 level in CR-fed female mice relative to AL-fed controls (**Fig. 1A**).

To determine the role of FGF21 in the metabolic response to a CR diet, we placed wild type (WT) and *Fgf21*^*-/-*^(KO) male and female mice on either *ad libitum* (AL) or calorie restricted (CR) diets for 15 weeks (**Figs. 1B, 2A**). We verified FGF21 is knocked-out using ELISA and examining *Fgf21* gene expression in the inguinal white adipose tissue (**Fig. S1A-D**). We utilized a 30% level of restriction using Engivo Global 2018 chow, a regimen which we and others have shown to extend lifespan in C57BL/6J mice (Mitchell et al., 2016; Pak et al., 2021). In contrast to our initial findings in CR (**Fig. 1A**), we recently showed that *Fgf21* has sex-dependent roles in the response of C57BL/6J mice to protein restriction (PR), with *Fgf21* being vital to many of the metabolic responses to PR in male but not in female mice (Green, Pak, et al., 2022). We therefore hypothesized that loss of *Fgf21* might have sex-dependent impacts on the response to CR, particularly with respect to metabolic health improvements.

CR is well known to reduce weight gain and adiposity, and we therefore followed the weight and body composition of the mice during the course of the study. Both male WT and male KO mice had reduced weight gain on a CR diet, with a reduction in both lean mass and fat mass gain; the overall effect, as we expected, was one of reduced adiposity (**Figs. 1C-J**). While the reduction in weight and lean mass by CR was very similar in both WT and KO mice, there was a greater reduction of fat mass and adiposity in WT mice due to a reduced level of fat and reduced adiposity in AL-fed KO mice. In contrast, in female mice a CR diet reduced weight gain and lean mass gain in both WT and KO mice, but did not have decrease fat mass gain or adiposity in either genotype (**Figs. 2B-I**). In accordance with the similar response of both WT and KO mice, food intake was similar between WT and KO mice in both sexes (**Figs. 1K-N, 2J-M**).

**Figure 2.**
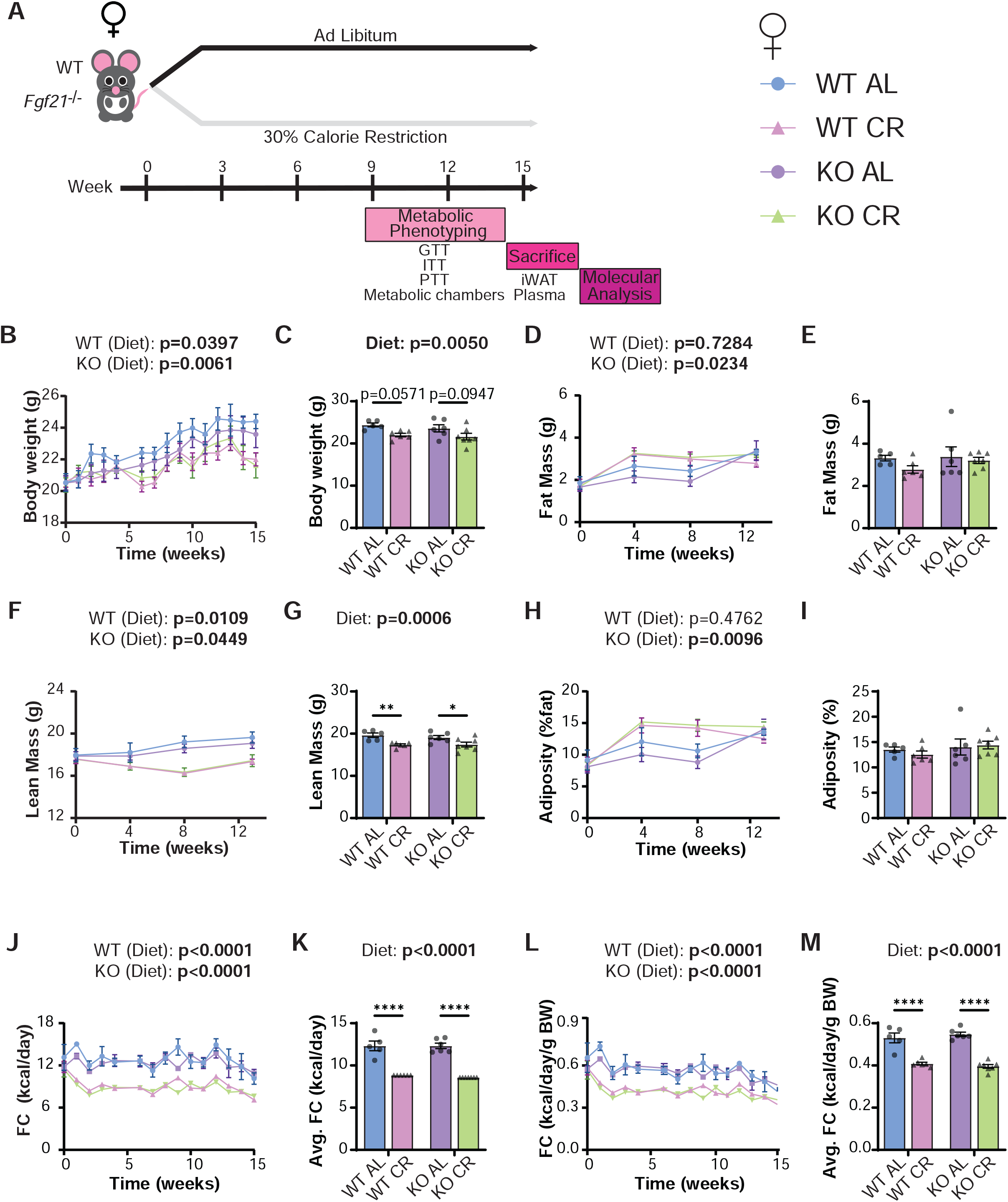
Loss of *Fgf21* does not impact the effect of CR on body weight, body composition and food consumption in female mice. (A) Experimental design. (B-C) Body weight measurement of female mice on the indicated diets (B) with final body weight at 15 weeks (C). (D-E) Fat mass measurement of female mice on the indicated diets (D) with final fat mass at 13 weeks (E). (F-G) Lean mass measurement of female mice on the indicated diets (F) with final lean mass at 13 weeks (G). (H-I) Adiposity of female mice on the indicated diets (H) with final adiposity at 13 weeks (I). (J-K) Food consumption (kcal/day) of female mice on the indicated diets (J) with average food consumed (K). (L-M) Food consumption per gram of body weight (kcal/day/g BW) of female mice on the indicated diets (L) with average food consumed per gram of body weight (M). (B-M) n=5-7 mice/group. For longitudinal studies, statistics for the overall effects of genotype, diet, and the interaction represent the p value from a two-way RM ANOVA or residual maximum likelihood (REML) analysis conducted individually for each genotype. For analyses of weight or body composition as a single time point, or analysis of average food consumption, statistics for the overall effects of genotype, diet, and the interaction represent the p value from a two-way ANOVA; *p<0.05, **p<0.01, **p<0.001, ****p<0.0001 Sidak’s test post 2-way ANOVA. Data represented as mean ± SEM.

### Loss of *Fgf21* has sex-specific impacts on aspects of the response of glucose homeostasis to CR

In mammals fed a CR diet, one of the most striking and broadly conserved effects is improved glucose homeostasis, which manifests in improved glucose tolerance and improved sensitivity to insulin. As administration of FGF21 promotes insulin sensitivity, we hypothesized that mice lacking *Fgf21* would have a reduced improvement in glucose homeostasis when fed a CR diet. After 8 weeks, which we and others have found is sufficient for CR to improve glucose homeostasis (Pak et al., 2021; Solon-Biet et al., 2015), we performed glucose, insulin, and pyruvate tolerance tests. We performed GTTs and ITTs after both a 7- and 21-hour fasting period to better understand their post-prandial glucose regulation (7 hours) and glucose regulation after a fasting period in which CR animals are accustomed to (21 hours). Although we understand that typically animals fast for 3-5 hours to prevent drastic changes in liver function, performing these tests at these times may provide better insight into the regulation of glucose homeostasis in a CR-fed mouse.

We observed that both male WT and KO mice had robust improvements in glucose tolerance when fed a CR diet, whether fasted for 7 hours or 21 hours (**Figs. 3A-B**). In agreement with other recent work from our lab, while CR-fed mice of both genotypes were extremely insulin sensitive following a prolonged fast, both WT and KO CR-fed mice had a post-prandial increase in insulin resistance, which is suggestive of having a tighter regulation of their blood glucose levels (**Fig. 3C**). After a longer fast, CR-fed mice regardless of genotype were more insulin sensitive (**Fig. 3D**). Fasting glucose and insulin levels were similar in all groups of male mice (**Fig. 3E-H**). However, KO CR-fed males had increased fasting blood glucose compared to WT CR-fed males after a 21 hour fast (**Figs. 3F**). Additionally, after refeeding, KO AL-fed males had higher insulin levels than mice in any other group (**Fig. 3H**).

**Figure 3.**
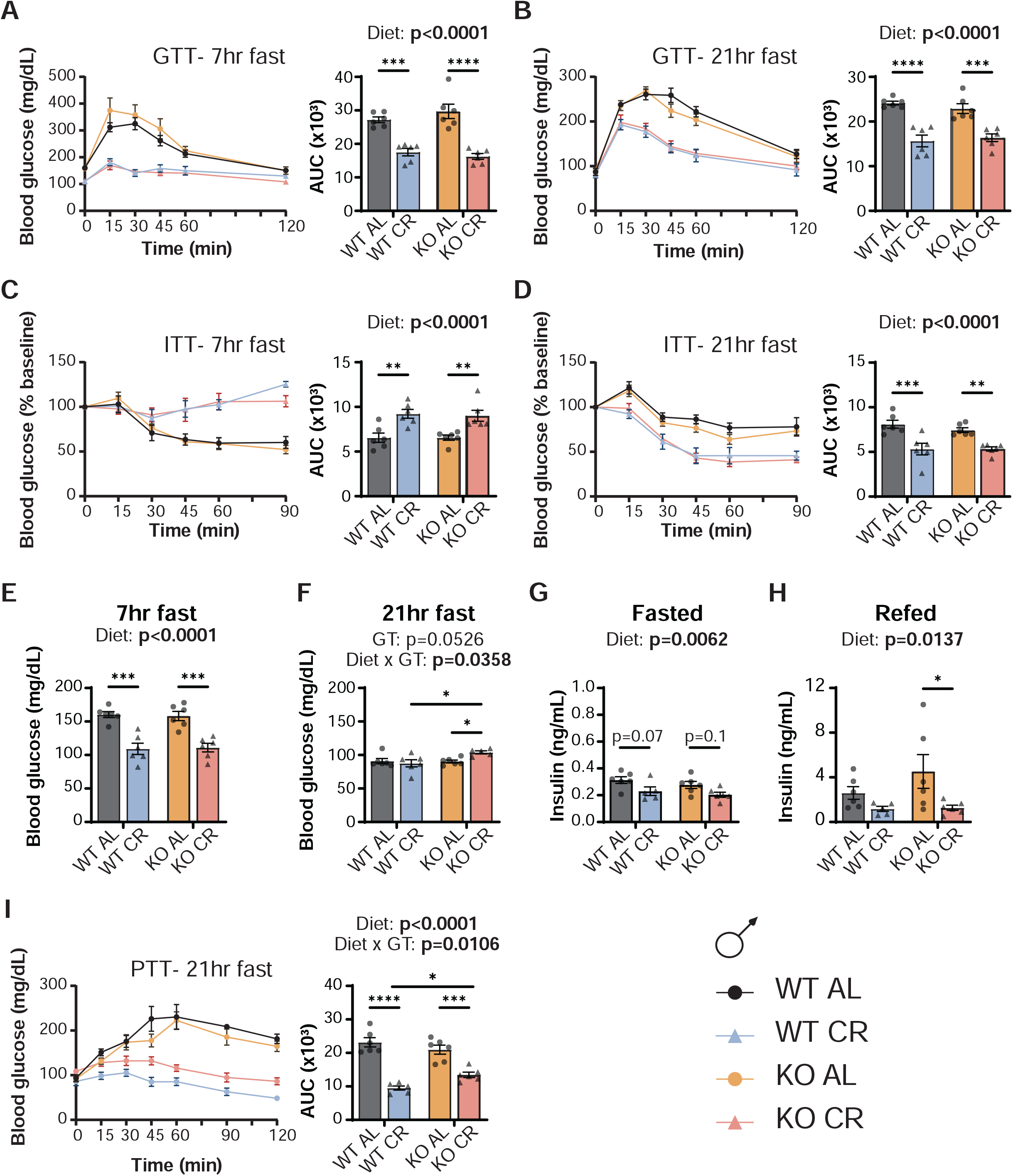
FGF21 is largely dispensable for the effects of CR on glucose homeostasis in male mice. (A-B) A glucose tolerance test (GTT) was conducted after a 7-hour (A) or 21-hour (B) fast. (C-D) An insulin tolerance test (ITT) was conducted after a 7-hour (C) or 21-hour (D) fast. (E-F) Fasting blood glucose (FBG) was measured after a 7-hour (E) or 21-hour (F) fast prior to the start of the ITT in panels C & D. (G-H) Plasma insulin was determined after a 16-hour fast (G) and after a 3-hour refeeding (H) after 15 weeks on diet. (I) A pyruvate tolerance test (PTT) was conducted after a 21-hour fast. (A-F, I) GTTs, ITTs, and PTT were performed between 8-13 weeks on diet regimens. (A-I) n=5-6 mice/group; statistics for the overall effects of genotype, diet, and the interaction represent the p value from a two-way ANOVA, *p<0.05, **p<0.01, **p<0.001, ****p<0.0001 from a Sidak’s post-test examining the effect of parameters identified as significant in the 2-way ANOVA. Data represented as mean ± SEM.

Investigating the mechanisms by which CR improves glucose tolerance, we found that CR-fed mice of both genotypes had significantly improved tolerance to pyruvate, indicating improved suppression of hepatic gluconeogenesis (**Fig. 3I**). However, CR-fed WT male mice were more pyruvate tolerant than CR-fed KO mice (**Fig. 3I**).

In agreement with what we observed in male mice, we observed improved glucose tolerance with CR in both WT and KO females after 7 hours of fasting (**Fig. 4A**). However, the effect of CR after 21 hours of fasting was significantly different only in WT mice; CR did not significantly improve glucose tolerance in KO females at this timepoint (**Fig. 4B**). Similar to males, CR-fed females of both genotypes were extremely insulin sensitive following a prolonged fast; however, unlike in males, KO CR-fed mice did not have a post-prandial increase in insulin resistance (**Figs. 4C-D**). Fasting insulin and glucose levels in KO female mice responded to CR differently than KO male mice (**Fig. 4E-H**). KO CR-fed females had increased fasting blood glucose compared to WT CR-fed females after 16 or 21 hours of fasting, while males only showed this effect after 21 hours of fasting (**Figs. 4E-F**). With regards to fasting insulin levels, female mice did not show a statistically significant effect of diet, and KO CR-fed females had significantly higher insulin levels following refeeding than their WT CR-fed counterparts (**Fig. 4G-H**). As with males, CR-fed females of both genotypes show improved suppression of hepatic gluconeogenesis during a pyruvate tolerance test (**Fig. 4I**). We conclude that *Fgf21* plays a role in the regulation of glucose metabolism in CR-fed female mice, but not in male mice.

**Figure 4.**
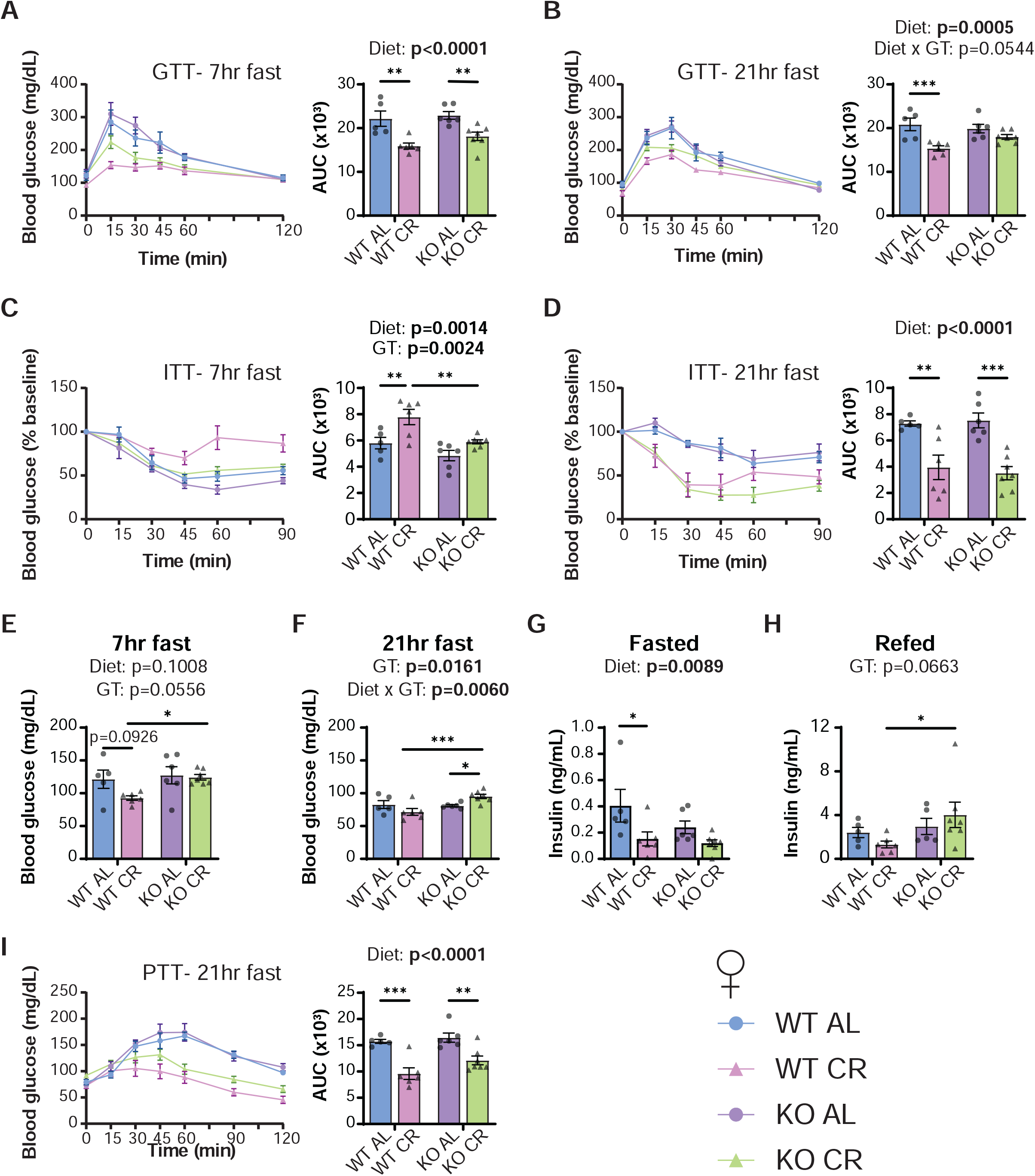
Loss of *Fgf21* impacts glucose homeostasis in female mice under specific feeding conditions. (A-B) A glucose tolerance test (GTT) was conducted after a 7-hour (A) or 21-hour (B) fast. (C-D) An insulin tolerance test (ITT) was conducted after a 7-hour (C) or 21-hour (D) fast. (E-F) Fasting blood glucose (FBG) was measured after a 7-hour (E) or 21-hour (F) fast prior to the start of the ITT in panels C & D. (G-H) Plasma insulin was determined after a 16-hour fast (G) and after a 3-hour refeeding (H) after 15 weeks on diet. (I) A pyruvate tolerance test (PTT) was conducted after a 21-hour fast. (A-F, I) GTTs, ITTs, and PTT were performed between 8-13 weeks on diet regimens. (A-I) n=5-7 mice/group; statistics for the overall effects of genotype, diet, and the interaction represent the p value from a two-way ANOVA, *p<0.05, **p<0.01, **p<0.001, ****p<0.0001 from a Sidak’s post-test examining the effect of parameters identified as significant in the 2-way ANOVA. Data represented as mean ± SEM.

### Loss of *Fgf21* does not affect CR-induced changes in components of energy balance and fuel utilization

We next observed respiration, energy expenditure and spontaneous activity using metabolic CLAMS-HC cages. CR has been previously shown to have a very distinct RER curve and reduced energy expenditure (Pak et al., 2021). As *Fgf21* is a critical regulator of energy balance through modulation of food intake and energy expenditure, we hypothesized that loss of *Fgf21* will affect energy balance in CR-fed mice. We therefore placed the mice in metabolic chambers, allowing us to determine energy expenditure via indirect calorimetry while also assessing food consumption, activity, and fuel source utilization.

We examined the contribution of diet and genotype to energy expenditure in male mice over a 24-hour period, correcting for differences in body weight (**Fig. 5A**) and lean mass (**Fig. 5B**) using analysis of covariance (ANCOVA). ANCOVA analysis assumes a linear relationship between the variables and their covariates. If the slope is equal between groups, then the regression lines are parallel, and elevation is then tested to determine any differences (i.e., if slopes are statistically significantly different, elevation will not be determined). Male mice on CR diets had decreased energy expenditure relative to AL-fed male mice, regardless of genotype (**Figs. 5A-B**). CR mice undergo rapid lipogenesis following refeeding, then sustain themselves via the utilization of these stored lipids (Bruss, Khambatta, Ruby, Aggarwal, & Hellerstein, 2010; Yu et al., 2019). We determined substrate utilization by examining the respiratory exchange ratio (RER), the ratio of O_2_ consumed and CO_2_ produced; a value close to 1.0 indicates that carbohydrates are primarily utilized for energy production, and for active *de novo* lipogenesis while a value approaching 0.7 indicates that lipids are the predominant energy source (Hasek et al., 2010). Both WT and KO CR-fed male mice displayed a distinct pattern in RER, with a rapid increase in RER indicating lipogenesis following once-per-day feeding (**Fig. 5C-D**). We also observed no effect of genotype on spontaneous activity, although there was an overall trend of CR-fed males towards increased activity (p=0.06) (**Fig. 5E**). Female mice behaved similarly to male mice, with strong effects of diet but not genotype on energy expenditure and RER (**Figs. 5F-I**). While female CR-fed mice also had an overall trend towards increased activity (p=0.0589), we observed a trend toward decreased activity of KO females (p=0.0719) (**Figs. 5J**).

**Figure 5.**
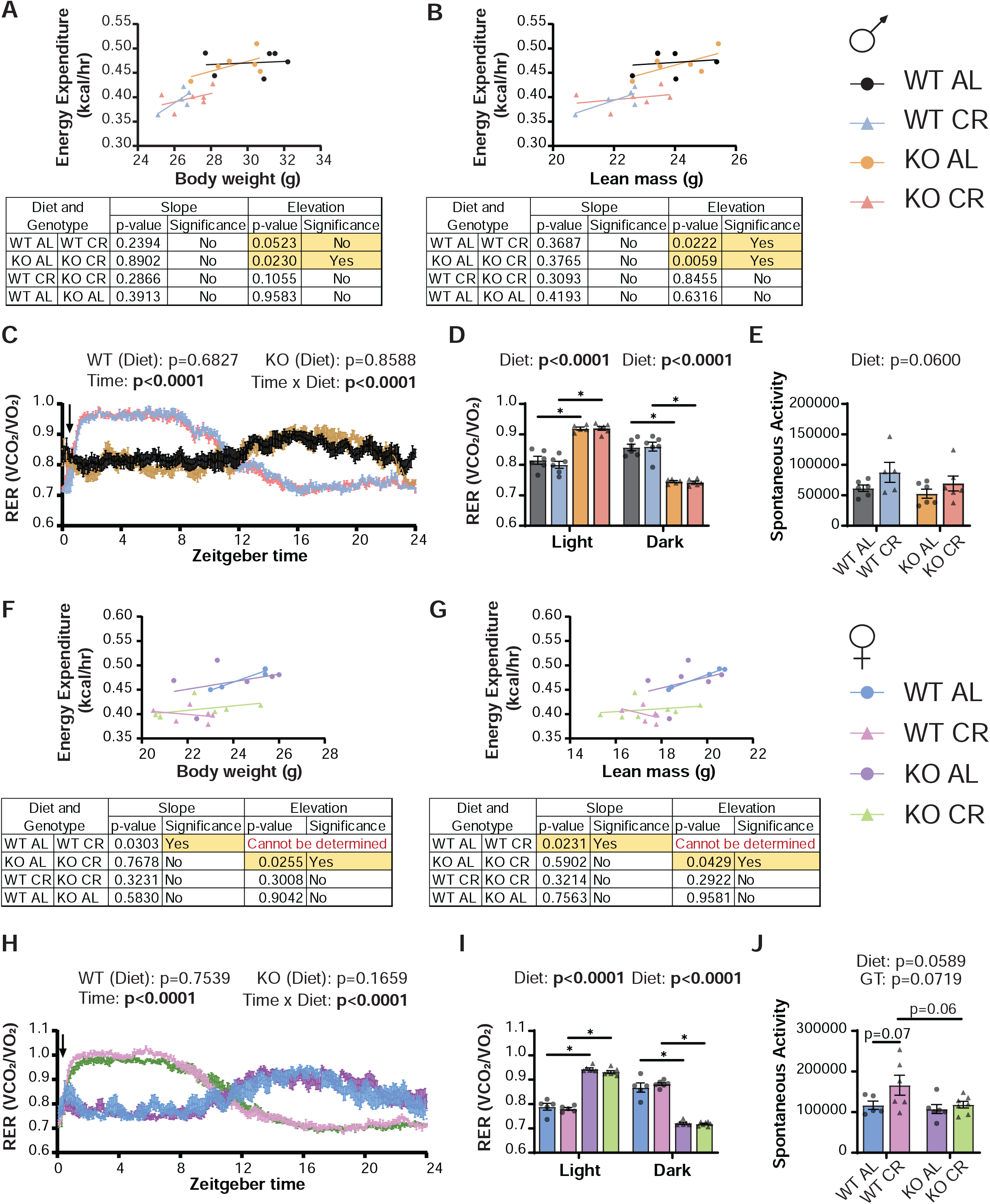
Loss of *Fgf21* does not affect heat, respiratory exchange ratio (RER) or spontaneous activity in CR-fed male and female mice. (A-B) Energy expenditure of male mice as a function of body weight (A) and lean mass (B) over a 24-hour period. (C-D) Respiratory Exchange Ratio (RER) of male mice over the course of a 24-hour period (C) or averaged during the light and dark cycles (D). Arrow indicates feeding time of CR mice. (E) Spontaneous activity of male mice. (F-G) Energy expenditure of female mice as a function of body weight (F) and lean mass (G) over a 24-hour period. (H-I) RER of female mice over the course of a 24-hour period (H) or averaged during the light and dark cycles (I). Arrow indicates feeding time of CR mice. (J) Spontaneous activity of female mice. (A-J) Energy expenditure, RER and spontaneous activity was determined after mice were on the dietary regimens for 13-15 weeks. n=5-6 (males), n=5-7 (females). (A-B, F-G) data for each individual mouse is plotted; simple linear regression (ANCOVA) was calculated to determine if the slopes or elevations are equal; if the slopes are significantly different, differences in elevation cannot be determined. (C, H) statistics for the overall effects of genotype, diet, and the interaction represent the p value from a two-way RM ANOVA or REML analysis conducted individually for each genotype. (D, I) statistics for the overall effects of genotype, diet, and the interaction represent the p value from a two-way ANOVA conducted separately for the light and dark cycles; *p<0.05, **p<0.01, **p<0.001, ****p<0.0001 from a Sidak’s post-test examining the effect of parameters identified as significant in the 2-way ANOVA. (E, J) statistics for the overall effects of genotype, diet, and the interaction represent the p value from a two-way ANOVA; *p<0.05, Sidak’s test post 2-way ANOVA. Data represented as mean ± SEM.

### Loss of *Fgf21* blunts CR-induced reprogramming of white adipose tissues in males but not females

Adipose tissue plays a central role in the response to CR (Miller et al., 2017); in mice, CR promotes the beiging of iWAT, increasing the expression of thermogenic, lipogenic, and lipolytic genes (Sheng et al., 2021; Yu et al., 2019). FGF21 plays a key role in adipose tissue, promoting energy expenditure in response to various stimuli, in part through promoting the beiging of iWAT (Cuevas-Ramos, Mehta, & Aguilar-Salinas, 2019; Douris et al., 2015; Green, Lamming, et al., 2022; Hill et al., 2019; Klein Hazebroek & Keipert, 2022; Veniant et al., 2012). We therefore examined the role of FGF21 in CR-induced beiging of iWAT. We examined the expression of thermogenic, lipogenic and lipolytic gene expression, increases in which are characteristic of beiging iWAT as well as examined and quantified the iWAT histology.

As expected, in male CR-fed mice, expression of thermogenic genes *Ucp1, Cidea* and *Elovl3* were increased (**Figs. 6A-C**). For all three genes, there was a diminished effect of CR in the KO mice lacking *Fgf21*. We also looked at the lipogenic genes *Dgat1, Fasn* and *Acc1*. We found that a CR diet induced *Acc1* and *Dgat1* in both WT males and KO males; however, there was a significant effect of genotype and a significant diet x genotype interaction on the induction of *Fasn* by CR, with significantly less induction of *Fasn* in KO than in WT males (**Figs. 6D-F**). Finally, we examined the expression of two lipolytic genes, *Atgl* and *Lipe*. Similar to *Fasn*, we found a significant induction of both genes by CR in WT males, but a significant effect of genotype and a significant diet x genotype interaction resulting from a blunting of the induction of these genes in CR-fed KO males (**Figs. 6G-H**). Finally, we characterized iWAT histology. We observed a strong visual effect of CR on multilocularity in WT males, indicative of adipose tissue beiging, but a reduced effect in KO mice, which is supported by the quantification of multilocular cells per animal (**Fig. 6I-K**). Therefore, loss of Fgf21 seems to slightly blunt CR-induced beiging of male mice.

**Figure 6:**
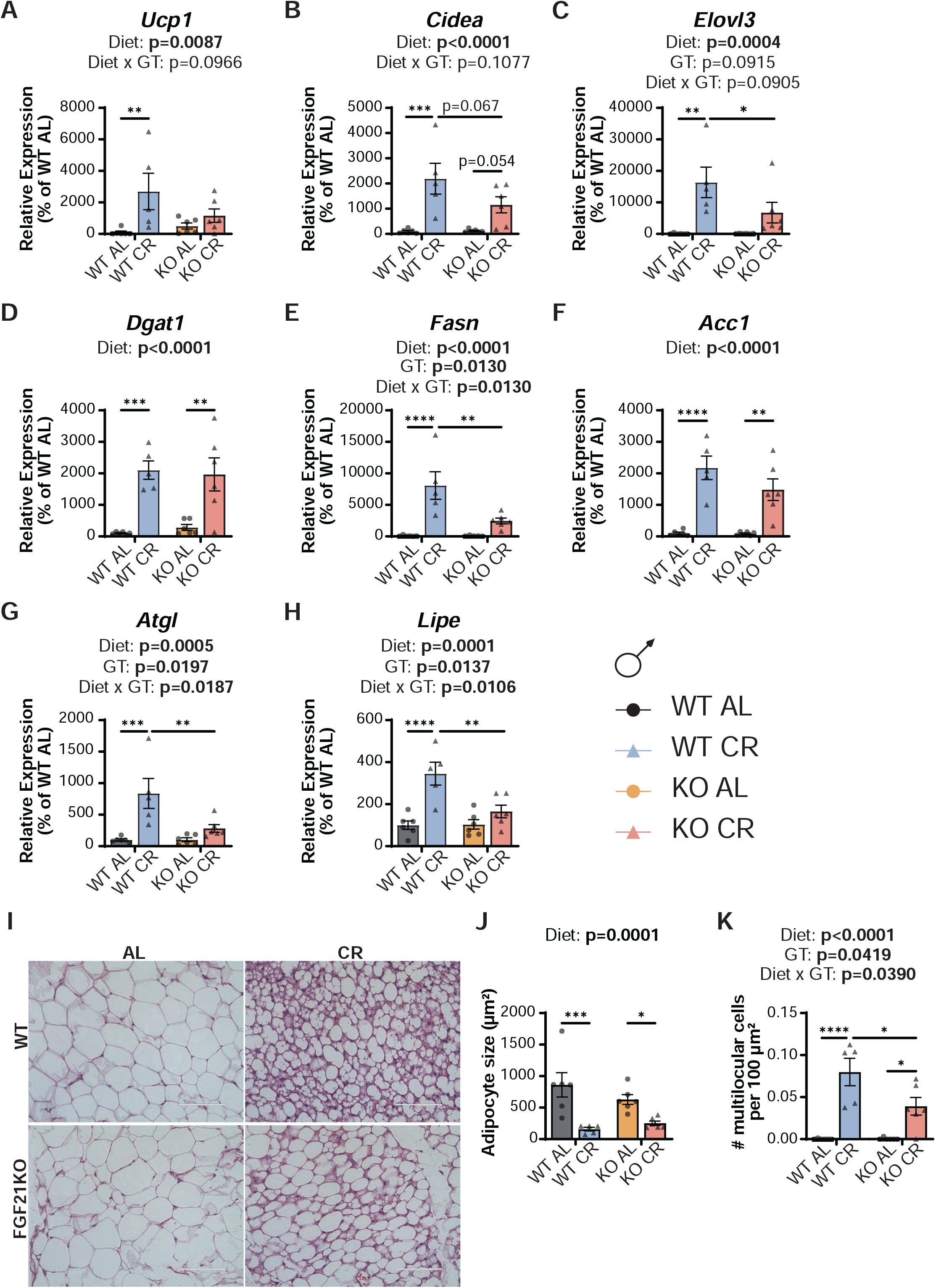
Loss of *Fgf21* blunts CR-induced beiging in the inguinal white adipose tissue of male mice. (A-C) The expression of three thermogenic genes, *Ucp1* (A), *Cidea* (B) and *Elovl3* (C) was quantified in the inguinal white adipose tissue (iWAT) of male mice. (D-F) The expression of three lipogenic genes, *Dgat1* (D), *Fasn* (E) and *Acc1* (F) was quantified in the iWAT of male mice. (G-H) The expression of the lipolytic genes *Atgl* (G) and *Lipe* (H) was quantified in the iWAT of male mice. (I) Hematoxylin and eosin (HE) staining (representative images; scale bar=100µm, 40X magnification) from iWAT of male mice. (J) Quantified adipocyte size (µm^2^) from HE-stained iWAT images from male mice. (K) Number of multilocular cells per 100 µm^2^ of HE-stained iWAT images of male mice. (A-H, J-K) n=5-6 mice/group; statistics for the overall effects of genotype, diet, and the interaction represent the p value from a two-way ANOVA, *p < 0.05, **p<0.01, **p<0.001, ****p<0.0001 from a Sidak’s post-test examining the effect of parameters identified as significant in the 2-way ANOVA. Data represented as mean ± SEM.

In contrast, in female mice we observed a substantially decreased effect of CR on gene expression as compared to males, even in WT mice (**Figs. 7A-H**). There was not a significant effect of CR on the expression of either *Ucp1* or *Lipe*. There was a significant overall effect of CR on the expression of the thermogenic genes *Cidea* and *Elovl3*, but in contrast to males a similar effect was seen in both WT and KO females (**Fig. 7B-C**). Also, in contrast to males, deletion of *Fgf21* did not impact lipogenic or lipolytic gene expression in CR-fed females (**Fig. 7D-H**). Histological examination of iWAT suggests that CR increases beiging equally well in both WT and KO female mice (**Fig. 7I-K**).

**Figure 7:**
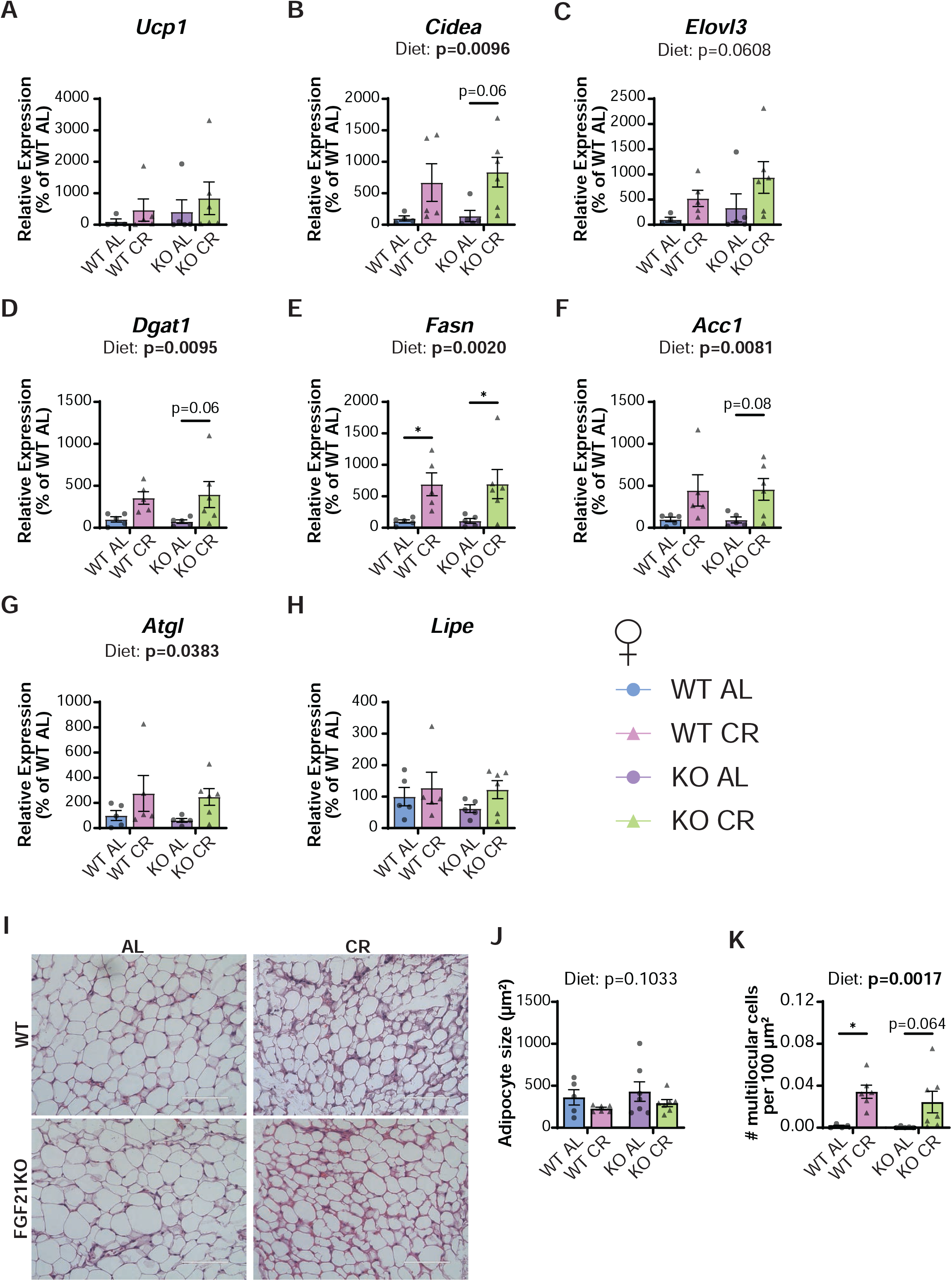
Loss of *Fgf21* does not impact CR-induced beiging in the inguinal white adipose tissue of female mice. (A-C) The expression of three thermogenic genes, *Ucp1* (A), *Cidea* (B) and *Elovl3* (C) was quantified in the inguinal white adipose tissue (iWAT) of female mice. (D-F) The expression of three lipogenic genes, *Dgat1* (D), *Fasn* (E) and *Acc1* (F), was quantified in the iWAT of female mice. (G-H) The expression of the lipolytic genes *Atgl* (G) and *Lipe* (H) was quantified in the iWAT of female mice. (I) Hematoxylin and eosin (HE) staining (representative images; scale bar = 100µm, 40X magnification) from iWAT of female mice. (J) Quantified adipocyte size (µm^2^) from HE-stained iWAT images from female mice. (K) Number of multilocular cells per 100 µm^2^ of HE-stained iWAT images of female mice. (A-H, J-K) n=4-7 mice/group; statistics for the overall effects of genotype, diet, and the interaction represent the p value from a two-way ANOVA, *p < 0.05, **p<0.01, **p<0.001, ****p<0.0001 from a Sidak’s post-test examining the effect of parameters identified as significant in the 2-way ANOVA. Data represented as mean ± SEM.

## Discussion

Calorie restriction improves metabolic health and longevity across diverse species (Belsky, Huffman, Pieper, Shalev, & Kraus, 2017; Kraus et al., 2019; Mattison et al., 2017; Pak et al., 2021; Rhoads et al., 2020). Many of the metabolic effects of CR are at first glance similar to those induced by the energy balance hormone FGF21. FGF21 promotes the browning/beiging of iWAT as well as activating BAT, thereby leading to increased energy expenditure and reduced body weight, and overexpression of FGF21 is sufficient to extend mouse lifespan (Cuevas-Ramos et al., 2019; Douris et al., 2015; Green, Lamming, et al., 2022; Veniant et al., 2012; Zhang et al., 2012). Some work has identified a role for FGF21 in specific phenotypes of CR in male mice and rats (Fujii et al., 2019; Thompson et al., 2014), and we found that morning-fed CR mice display a sex-specific induction of FGF21 in their natural feeding state (**Figure S1**). While it is plausible that CR promotes metabolic health in part via FGF21, the requirement for FGF21 in the metabolic response to CR has not been comprehensively investigated.

Here, we examined the role of FGF21 in the metabolic response to 30% CR in male and female mice fed once per day in the morning, a paradigm widely used in previous studies. We find that deletion of *Fgf21* does not substantially alter the effect of CR on body weight, composition, or food consumption in either males or females (**Fig. 8A**). However, when we examined the regulation of glucose homeostasis, we identified several roles for FGF21 in the response to CR. In males, WT mice fed a CR diet had better suppression of glucose production from pyruvate and higher fasting blood glucose than WT CR-fed males. We observed a similar trend in females, with KO females also showing a blunted improvement in glucose tolerance on a CR diet as compared to WT mice when examined after a prolonged fast, and KO CR-fed females also had increased fasting blood glucose at two time points. WT female mice, but not KO females, showed a post-prandial increase in insulin resistance. Our results are consistent with a minor role for *Fgf21* in the suppression of hepatic glucose output during CR in both sexes, as well as literature showing that administration of FGF21 promotes hepatic insulin sensitivity (Gong et al., 2016).

**Figure 8:**
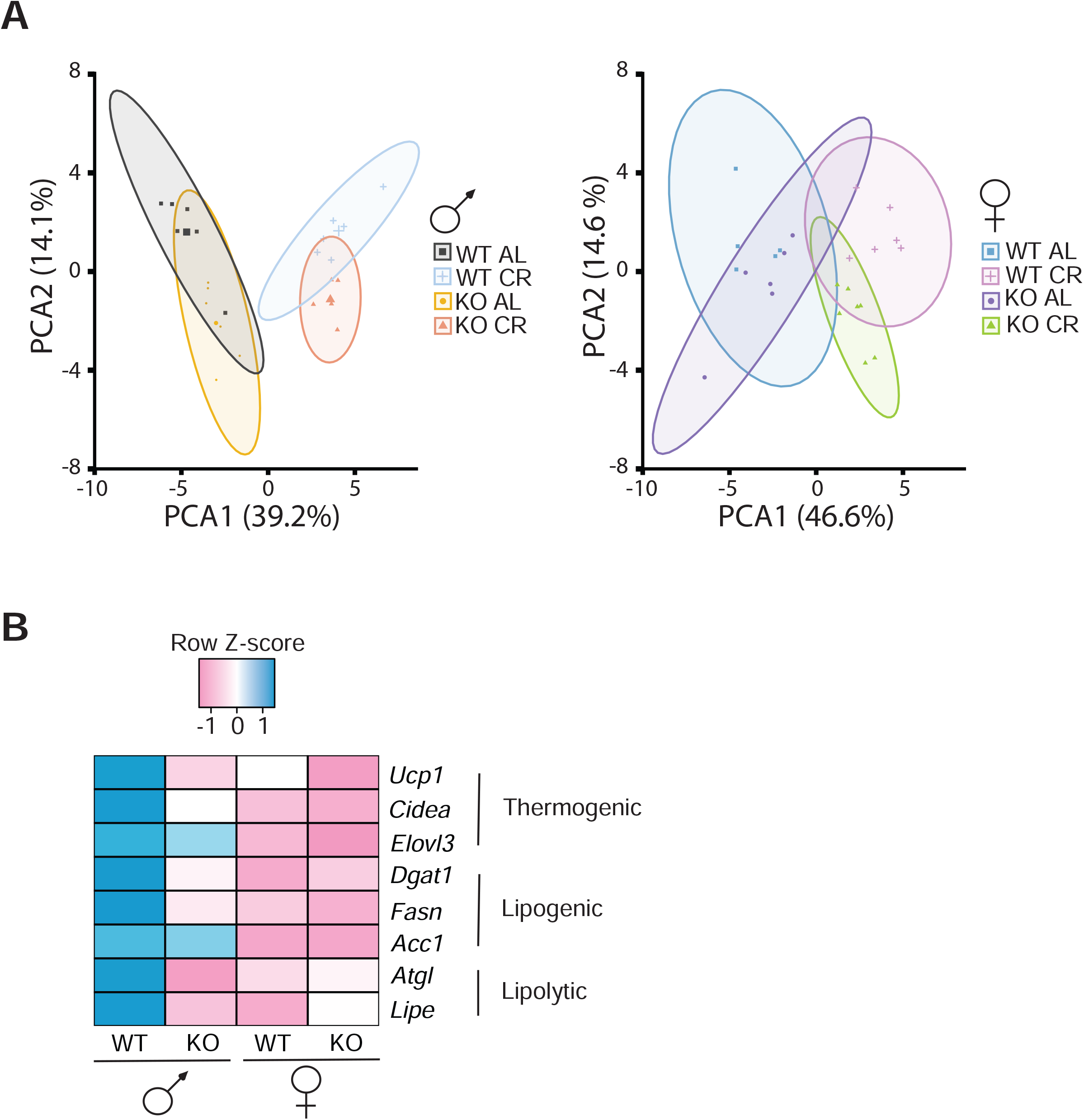
Loss of *Fgf21* does not affect the overall phenotypic response to CR, but blunts the induction of iWAT genes in a sex-specific manner. (A) Phenotypic measurements of individual mice visualized by genotype and diet groups and split by sex indicating separation along PC1 and PC2; 95% confidence ellipses are indicated. (B) Log_2_ fold changes in gene expression induced by CR in the iWAT of each genotype and sex of mice were z-scaled normalized.

Loss of *Fgf21* also does not substantially alter the effects of CR on energy balance in either male or female mice. This was rather surprising, as the role of FGF21 in these phenotypes in response to restriction of protein or dietary amino acids has been well established (Hill et al., 2017; Laeger et al., 2014; Wanders et al., 2017; Yu et al., 2021). A limitation of the literature is that much of this work has been performed in C57BL/6J male mice, and we have found that many effects of protein restriction depend on sex and strain. This includes the role of FGF21, which we have found plays an important role in the response to protein restriction in C57BL/6J males, but not C57BL/6J females (Green, Pak, et al., 2022). Our results are consistent with a model in which FGF21 is not essential for the effects of CR on energy balance.

In light of this conclusion, it is somewhat surprising that loss of *Fgf21* blunts the beiging of white adipose tissue of male mice placed on a CR diet (**Fig. 8B**). We find that CR-fed KO male mice do not have increased expression of the thermogenic gene *Ucp1*, typically thought of as a main driver for thermogenesis during beiging. However, the expression of two other thermogenic genes, *Cidea* and *Elovl3*, are also elevated by CR in males, although the induction of these genes by CR is blunted in the absence of FGF21. *Elovl3*, which leads to an increase in very long-chain fatty acids, is an adaptive thermogenesis marker found to be increased in brown adipose tissue in response to cold exposure (Fischer et al., 2020). Futile cycling between lipogenic and lipolytic gene expression has also been shown to be a characteristic of beiging white adipose tissue (Mottillo et al., 2014; Yu et al., 2021). We also found blunted expression of lipogenic and lipolytic genes in KO CR-fed male mice relative to WT CR-fed males. We also found a reduced effect of multilocular adipocytes at the histological level, suggesting that loss of *Fgf21* is required for the full induction of beiging by CR in male mice. Intriguingly, these effects are sex-specific, as in CR-fed female mice, beiging is not induced to the same extent as in the males, loss of *Fgf21* does not impact thermogenic, lipogenic or lipolytic gene induction in response to CR in females, and KO female mice on a CR diet have histologically similar iWAT to WT CR-fed female mice.

Limitations of this study include that our study lasted less than 4 months; there may be more prominent differences in loss of *Fgf21* under a longer CR duration that we did not see in this short time frame, and we did not examine the contribution of FGF21 to the effects of CR on frailty, cognition or lifespan. We examined the role of FGF21 on the response of iWAT to CR, but we did not examine the molecular response of other adipose depots or other tissues. We collected tissues for molecular examination in the refed state, but due to the distinct eating patterns of CR-fed mice, examining tissues in other feeding states might prove insightful. We fed CR mice in the morning shortly after lights on; while this intervention extends lifespan (Pak et al., 2021; Yu et al., 2019), recently it has been suggested that there may be distinct effects of feeding mice in the morning or evening (Acosta-Rodriguez et al., 2022), and examining CR mice fed in the evening might find different results. Finally, Green and colleagues showed that only C57BL/6J males had a robust induction of FGF21 in response to protein restriction (PR), while females had a reduced response (p=0.056) and FGF21 did not increase in response to PR in either sex of DBA/2J or UM-HET3 mice (Green, Pak, et al., 2022). Examining the response to CR in multiple strains might reveal more details about the role of FGF21 as well as FGF21-independent mechanisms in the metabolic response to CR.

In summary, here we have tested the hypothesis that the nutrient responsive hormone FGF21 is a key mediator of the CR-induced improvements in body composition, glucose homeostasis, energy balance. We have determined that deletion of *Fgf21* from the whole body does not impair the ability of CR to improve body composition or energy balance and has relatively minor effects on CR-induced improvements in glucose regulation. In agreement with the literature linking FGF21 to the beiging of iWAT, we find that FGF21 plays a key role in CR-induced beiging of iWAT, but only in males. Here, we disprove the hypothesis that FGF21 mediates the metabolic benefits of CR, bringing us one step closer to identifying the true molecular mediators of the beneficial effects of CR. Notably, we have not excluded a role for FGF21 in the effects of CR on frailty or lifespan. Additionally, we have not excluded the role of factors upstream of FGF21, including the general control nonderepressible 2 (GCN2) kinase, activating transcription factor 4 (ATF4), nor factors downstream of FGF21, such as insulin growth factor 1 (IGF1) and mechanistic target of rapamycin (mTOR), in the response to CR.

## ACKNOWLEDGEMENTS

We would like to thank all members of the Lamming lab, including those who provided any advice during this project. The Lamming laboratory is supported in part by the NIH/NIA (AG056771, AG062328, and AG061635 to D.W.L.), NIH/NIDDK (DK125859 to D.W.L.) and startup funds from the University of Wisconsin-Madison School of Medicine and Public Health and Department of Medicine to D.W.L. M.F.C. was supported in part by a Diana Jacobs Kalman/AFAR Scholarships for Research in the Biology of Aging. C.Y.-Y. was supported by a training grant (T32AG000213) and a NIA F32 postdoctoral fellowship (F32AG077916). R.B. is supported by a training grant (T32DK007665). C.L.G. was supported in part by a Glenn/AFAR postdoctoral fellowship. H.H.P. was supported by a NIA F31 predoctoral fellowship (AG066311). Support for this research was provided by the University of Wisconsin - Madison Office of the Vice Chancellor for Research and Graduate Education with funding from the Wisconsin Alumni Research Foundation. The UW Carbone Cancer Center (UWCCC) Experimental Pathology Laboratory is supported by P30 CA014520 from the NIH/NCI. This work was supported in part by the U.S. Department of Veterans Affairs (I01-BX004031), and this work was supported using facilities and resources from the William S. Middleton Memorial Veterans Hospital. The content is solely the responsibility of the authors and does not necessarily represent the official views of the NIH. This work does not represent the views of the Department of Veterans Affairs or the United States Government.

## Data Availability Statement

The data that support the findings of this study are available from the corresponding author upon reasonable request.

## AUTHOR CONTRIBUTIONS

MFC, HHP, and DWL conceived of and designed the experiments. MFC, IA, CYY, RB, AMB, IK, and MMS performed the experiments. MFC, IK, CLG and DWL analyzed the data and wrote the manuscript.

---

**Supplementary Figure 1:**
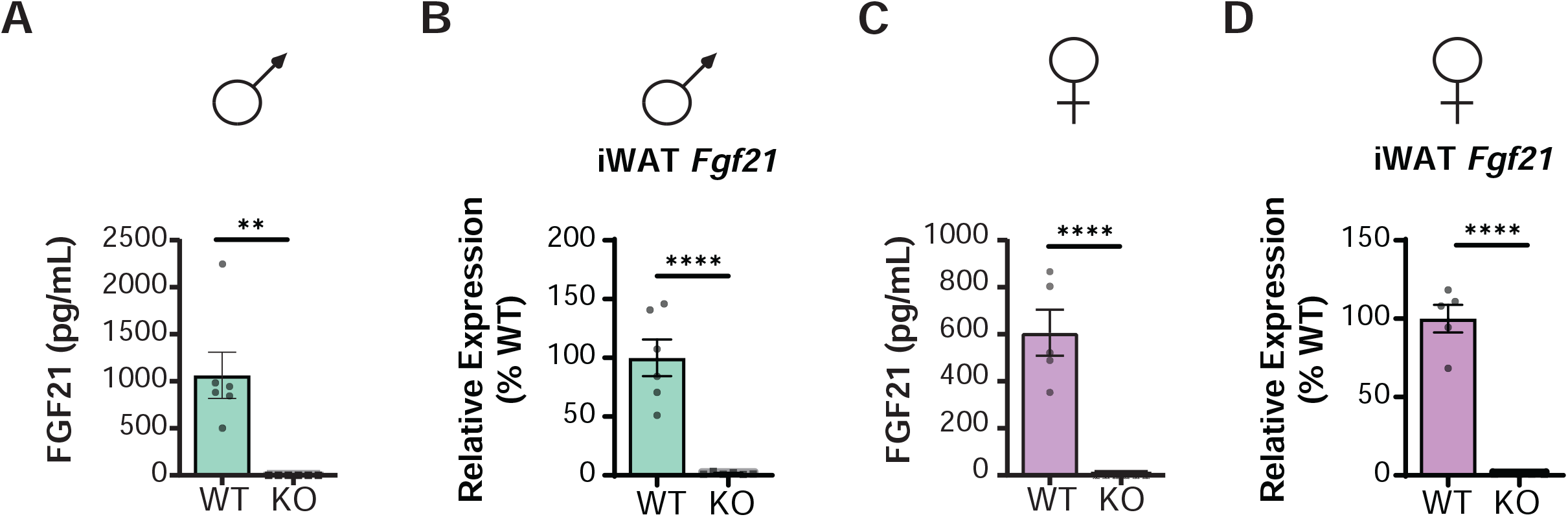
Confirmation of *Fgf21* knock-out in circulation and inguinal white adipose tissue. (A) Circulating FGF21 was measured via ELISA in male WT and KO mice. (B) *Fgf21* gene expression was measured in the iWAT of male WT and KO mice. (C) Circulating FGF21 was measured via ELISA in female WT and KO mice. (D) *Fgf21* gene expression was measured in the iWAT of female WT and KO mice. (A-B) n=6 mice/group, ** =p=0.0015, **** = p<0.0001, unpaired, two-tailed t-test. (C-D) n=5-6 mice/ group **** = p<0.0001, unpaired, two-tailed t-test. Data represented as mean ± SEM.

**Supplementary Table 1:**
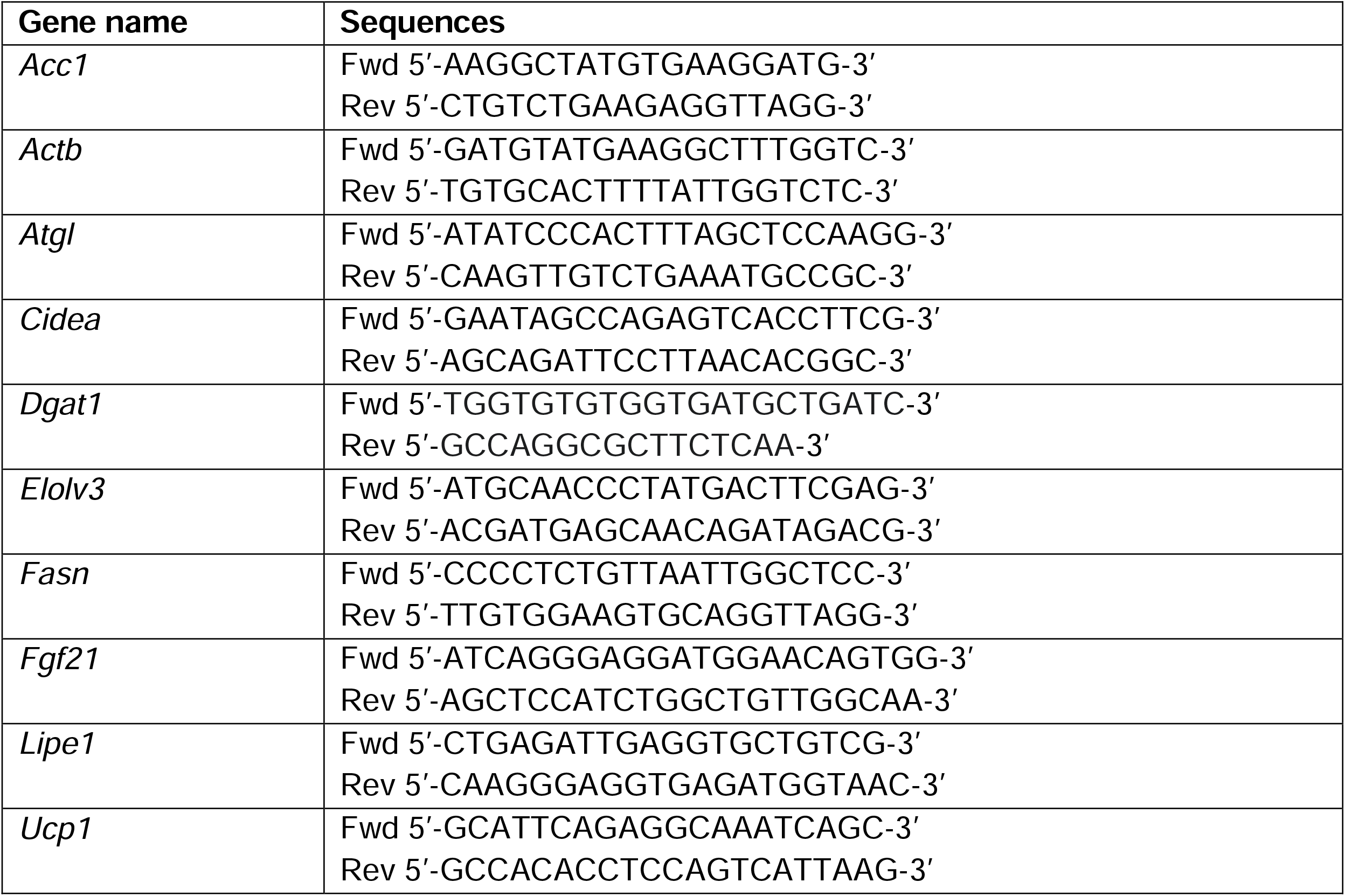
qPCR primer sequences.

## References

Acosta-Rodriguez, V., Rijo-Ferreira, F., Izumo, M., Xu, P., Wight-Carter, M., Green, C. B., & Takahashi, J. S. (2022). Circadian alignment of early onset caloric restriction promotes longevity in male C57BL/6J mice. Science, 376(6598), 1192–1202. doi:10.1126/science.abk0297

Balasubramanian, P., Schaar, A. E., Gustafson, G. E., Smith, A. B., Howell, P. R., Greenman, A., … Anderson, R. M. (2021). Adiponectin receptor agonist AdipoRon improves skeletal muscle function in aged mice. bioRxiv, 2021.2009.2016.460597. doi:10.1101/2021.09.16.460597

Bellantuono, I., de Cabo, R., Ehninger, D., Di Germanio, C., Lawrie, A., Miller, J., … Lamming, D. W. (2020). A toolbox for the longitudinal assessment of healthspan in aging mice. Nat Protoc, 15(2), 540–574. doi:10.1038/s41596-019-0256-1

Belsky, D. W., Huffman, K. M., Pieper, C. F., Shalev, I., & Kraus, W. E. (2017). Change in the Rate of Biological Aging in Response to Caloric Restriction: CALERIE Biobank Analysis. J Gerontol A Biol Sci Med Sci, 73(1), 4–10. doi:10.1093/gerona/glx096

Bonkowski, M. S., Rocha, J. S., Masternak, M. M., Al Regaiey, K. A., & Bartke, A. (2006). Targeted disruption of growth hormone receptor interferes with the beneficial actions of calorie restriction. Proc Natl Acad Sci U S A, 103(20), 7901–7905. doi:10.1073/pnas.0600161103

Bruss, M. D., Khambatta, C. F., Ruby, M. A., Aggarwal, I., & Hellerstein, M. K. (2010). Calorie restriction increases fatty acid synthesis and whole body fat oxidation rates. Am J Physiol Endocrinol Metab, 298(1), E108–116. doi:10.1152/ajpendo.00524.2009

Coskun, T., Bina, H. A., Schneider, M. A., Dunbar, J. D., Hu, C. C., Chen, Y., … Kharitonenkov, A. (2008). Fibroblast growth factor 21 corrects obesity in mice. Endocrinology, 149(12), 6018–6027. doi:10.1210/en.2008-0816

Cuevas-Ramos, D., Mehta, R., & Aguilar-Salinas, C. A. (2019). Fibroblast Growth Factor 21 and Browning of White Adipose Tissue. Front Physiol, 10, 37. doi:10.3389/fphys.2019.00037

Cummings, N. E., Williams, E. M., Kasza, I., Konon, E. N., Schaid, M. D., Schmidt, B. A., … Lamming, D. W. (2018). Restoration of metabolic health by decreased consumption of branched-chain amino acids. J Physiol, 596(4), 623–645. doi:10.1113/JP275075

Douris, N., Stevanovic, D. M., Fisher, F. M., Cisu, T. I., Chee, M. J., Nguyen, N. L., … Maratos-Flier, E. (2015). Central Fibroblast Growth Factor 21 Browns White Fat via Sympathetic Action in Male Mice. Endocrinology, 156(7), 2470–2481. doi:10.1210/en.2014-2001

Fischer, A. W., Behrens, J., Sass, F., Schlein, C., Heine, M., Pertzborn, P., … Heeren, J. (2020). Brown adipose tissue lipoprotein and glucose disposal is not determined by thermogenesis in uncoupling protein 1-deficient mice. J Lipid Res, 61(11), 1377–1389. doi:10.1194/jlr.RA119000455

Fontana, L., Cummings, N. E., Arriola Apelo, S. I., Neuman, J. C., Kasza, I., Schmidt, B. A., … Lamming, D. W. (2016). Decreased Consumption of Branched-Chain Amino Acids Improves Metabolic Health. Cell Rep, 16(2), 520–530. doi:10.1016/j.celrep.2016.05.092

Fujii, N., Uta, S., Kobayashi, M., Sato, T., Okita, N., & Higami, Y. (2019). Impact of aging and caloric restriction on fibroblast growth factor 21 signaling in rat white adipose tissue. Exp Gerontol, 118, 55–64. doi:10.1016/j.exger.2019.01.001

Gong, Q., Hu, Z., Zhang, F., Cui, A., Chen, X., Jiang, H., … Li, Y. (2016). Fibroblast growth factor 21 improves hepatic insulin sensitivity by inhibiting mammalian target of rapamycin complex 1 in mice. Hepatology, 64(2), 425–438. doi:10.1002/hep.28523

Green, C. L., Lamming, D. W., & Fontana, L. (2022). Molecular mechanisms of dietary restriction promoting health and longevity. Nat Rev Mol Cell Biol, 23(1), 56–73. doi:10.1038/s41580-021-00411-4

Green, C. L., Pak, H. H., Richardson, N. E., Flores, V., Yu, D., Tomasiewicz, J. L., … Lamming, D. W. (2022). Sex and genetic background define the metabolic, physiologic, and molecular response to protein restriction. Cell Metab, 34(2), 209–226 e205. doi:10.1016/j.cmet.2021.12.018

Hasek, B. E., Stewart, L. K., Henagan, T. M., Boudreau, A., Lenard, N. R., Black, C., … Gettys, T. W. (2010). Dietary methionine restriction enhances metabolic flexibility and increases uncoupled respiration in both fed and fasted states. Am J Physiol Regul Integr Comp Physiol, 299(3), R728–739. doi:10.1152/ajpregu.00837.2009

Hill, C. M., Laeger, T., Albarado, D. C., McDougal, D. H., Berthoud, H. R., Munzberg, H., & Morrison, C. D. (2017). Low protein-induced increases in FGF21 drive UCP1-dependent metabolic but not thermoregulatory endpoints. Sci Rep, 7(1), 8209. doi:10.1038/s41598-017-07498-w

Hill, C. M., Laeger, T., Dehner, M., Albarado, D. C., Clarke, B., Wanders, D., … Morrison, C. D. (2019). FGF21 Signals Protein Status to the Brain and Adaptively Regulates Food Choice and Metabolism. Cell Rep, 27(10), 2934–2947 e2933. doi:10.1016/j.celrep.2019.05.022

Husson, F., Josse, J., & Lê, S. (2008). FactoMineR: An R Package for Multivariate Analysis. Journal of Statistical Software, 25. doi:10.18637/jss.v025.i01

Inagaki, T., Dutchak, P., Zhao, G., Ding, X., Gautron, L., Parameswara, V., … Kliewer, S. A. (2007). Endocrine regulation of the fasting response by PPARalpha-mediated induction of fibroblast growth factor 21. Cell Metab, 5(6), 415–425. doi:10.1016/j.cmet.2007.05.003

Izumiya, Y., Bina, H. A., Ouchi, N., Akasaki, Y., Kharitonenkov, A., & Walsh, K. (2008). FGF21 is an Akt-regulated myokine. FEBS Lett, 582(27), 3805–3810. doi:10.1016/j.febslet.2008.10.021

Kassambara, A., & Mundt, F. (2020). factoextra: Extract and Visualize the Results of Multivariate Data Analyses. R package version 1.0.7. Retrieved from https://CRAN.R-project.org/package=factoextra

Klein Hazebroek, M., & Keipert, S. (2022). Obesity-resistance of UCP1-deficient mice associates with sustained FGF21 sensitivity in inguinal adipose tissue. Front Endocrinol (Lausanne), 13. doi:10.3389/fendo.2022.909621

Kraus, W. E., Bhapkar, M., Huffman, K. M., Pieper, C. F., Krupa Das, S., Redman, L. M., … Investigators, C. (2019). 2 years of calorie restriction and cardiometabolic risk (CALERIE): exploratory outcomes of a multicentre, phase 2, randomised controlled trial. Lancet Diabetes Endocrinol, 7(9), 673–683. doi:10.1016/S2213-8587(19)30151-2

Laeger, T., Henagan, T. M., Albarado, D. C., Redman, L. M., Bray, G. A., Noland, R. C., … Morrison, C. D. (2014). FGF21 is an endocrine signal of protein restriction. J Clin Invest, 124(9), 3913–3922. doi:10.1172/JCI74915

Lin, S. J., Defossez, P. A., & Guarente, L. (2000). Requirement of NAD and SIR2 for life-span extension by calorie restriction in Saccharomyces cerevisiae. Science, 289(5487), 2126–2128. doi:10.1126/science.289.5487.2126

Mattison, J. A., Colman, R. J., Beasley, T. M., Allison, D. B., Kemnitz, J. W., Roth, G. S., … Anderson, R. M. (2017). Caloric restriction improves health and survival of rhesus monkeys. Nat Commun, 8, 14063. doi:10.1038/ncomms14063

McCay, C. M., Crowell, M. F., & Maynard, L. A. (1935). The effect of retarded growth upon the length of life span and upon the ultimate body size. Journal of Nutrition, 10, 63–79.

Miller, K. N., Burhans, M. S., Clark, J. P., Howell, P. R., Polewski, M. A., DeMuth, T. M., … Anderson, R. M. (2017). Aging and caloric restriction impact adipose tissue, adiponectin, and circulating lipids. Aging Cell, 16(3), 497–507. doi:10.1111/acel.12575

Mitchell, S. J., Madrigal-Matute, J., Scheibye-Knudsen, M., Fang, E., Aon, M., Gonzalez-Reyes, J. A., … de Cabo, R. (2016). Effects of Sex, Strain, and Energy Intake on Hallmarks of Aging in Mice. Cell Metab, 23(6), 1093–1112. doi:10.1016/j.cmet.2016.05.027

Mottillo, E. P., Balasubramanian, P., Lee, Y. H., Weng, C., Kershaw, E. E., & Granneman, J. G. (2014). Coupling of lipolysis and de novo lipogenesis in brown, beige, and white adipose tissues during chronic beta3-adrenergic receptor activation. J Lipid Res, 55(11), 2276–2286. doi:10.1194/jlr.M050005

Nishimura, T., Nakatake, Y., Konishi, M., & Itoh, N. (2000). Identification of a novel FGF, FGF-21, preferentially expressed in the liver. Biochim Biophys Acta, 1492(1), 203–206. doi:10.1016/s0167-4781(00)00067-1

Pak, H. H., Haws, S. A., Green, C. L., Koller, M., Lavarias, M. T., Richardson, N. E., … Lamming, D. W. (2021). Fasting drives the metabolic, molecular and geroprotective effects of a calorie-restricted diet in mice. Nat Metab, 3(10), 1327–1341. doi:10.1038/s42255-021-00466-9

Potthoff, M. J., Inagaki, T., Satapati, S., Ding, X., He, T., Goetz, R., … Burgess, S. C. (2009). FGF21 induces PGC-1alpha and regulates carbohydrate and fatty acid metabolism during the adaptive starvation response. Proc Natl Acad Sci U S A, 106(26), 10853–10858. doi:10.1073/pnas.0904187106

Rhoads, T. W., Clark, J. P., Gustafson, G. E., Miller, K. N., Conklin, M. W., DeMuth, T. M., … Anderson, R. M. (2020). Molecular and Functional Networks Linked to Sarcopenia Prevention by Caloric Restriction in Rhesus Monkeys. Cell Syst, 10(2), 156–168 e155. doi:10.1016/j.cels.2019.12.002

Schwenk, F., Baron, U., & Rajewsky, K. (1995). A cre-transgenic mouse strain for the ubiquitous deletion of loxP-flanked gene segments including deletion in germ cells. Nucleic Acids Res, 23(24), 5080–5081. doi:10.1093/nar/23.24.5080

Sheng, Y., Xia, F., Chen, L., Lv, Y., Lv, S., Yu, J., … Ding, G. (2021). Differential Responses of White Adipose Tissue and Brown Adipose Tissue to Calorie Restriction During Aging. J Gerontol A Biol Sci Med Sci, 76(3), 393–399. doi:10.1093/gerona/glaa070

Solon-Biet, S. M., Mitchell, S. J., Coogan, S. C., Cogger, V. C., Gokarn, R., McMahon, A. C., … Le Couteur, D. G. (2015). Dietary Protein to Carbohydrate Ratio and Caloric Restriction: Comparing Metabolic Outcomes in Mice. Cell Rep, 11(10), 1529–1534. doi:10.1016/j.celrep.2015.05.007

Thompson, A. C., Bruss, M. D., Nag, N., Kharitonenkov, A., Adams, A. C., & Hellerstein, M. K. (2014). Fibroblast growth factor 21 is not required for the reductions in circulating insulin-like growth factor-1 or global cell proliferation rates in response to moderate calorie restriction in adult mice. PLoS ONE, 9(11), e111418. doi:10.1371/journal.pone.0111418

Turturro, A., Witt, W. W., Lewis, S., Hass, B. S., Lipman, R. D., & Hart, R. W. (1999). Growth curves and survival characteristics of the animals used in the Biomarkers of Aging Program. J Gerontol A Biol Sci Med Sci, 54(11), B492–501. doi:10.1093/gerona/54.11.b492

Veniant, M. M., Hale, C., Helmering, J., Chen, M. M., Stanislaus, S., Busby, J., … Lloyd, D. J. (2012). FGF21 promotes metabolic homeostasis via white adipose and leptin in mice. PLoS One, 7(7), e40164. doi:10.1371/journal.pone.0040164

Wanders, D., Forney, L. A., Stone, K. P., Burk, D. H., Pierse, A., & Gettys, T. W. (2017). FGF21 Mediates the Thermogenic and Insulin-Sensitizing Effects of Dietary Methionine Restriction but Not Its Effects on Hepatic Lipid Metabolism. Diabetes, 66(4), 858–867. doi:10.2337/db16-1212

Warnes, G., Bolker, B., Bonebakker, L., Gentleman, R., Huber, W., Liaw, A., … Möller, S. (2005). gplots: Various R programming tools for plotting data (Vol. 2).

Wei, S., Zhao, J., Bai, M., Li, C., Zhang, L., & Chen, Y. (2019). Comparison of glycemic improvement between intermittent calorie restriction and continuous calorie restriction in diabetic mice. Nutr Metab (Lond), 16, 60. doi:10.1186/s12986-019-0388-x

Xu, J., Stanislaus, S., Chinookoswong, N., Lau, Y. Y., Hager, T., Patel, J., … Veniant, M. M. (2009). Acute glucose-lowering and insulin-sensitizing action of FGF21 in insulin-resistant mouse models--association with liver and adipose tissue effects. Am J Physiol Endocrinol Metab, 297(5), E1105–1114. doi:10.1152/ajpendo.00348.2009

Yap, Y. W., Rusu, P. M., Chan, A. Y., Fam, B. C., Jungmann, A., Solon-Biet, S. M., … Rose, A. J. (2020). Restriction of essential amino acids dictates the systemic metabolic response to dietary protein dilution. Nat Commun, 11(1), 2894. doi:10.1038/s41467-020-16568-z

Yu, D., Richardson, N. E., Green, C. L., Spicer, A. B., Murphy, M. E., Flores, V., … Lamming, D. W. (2021). The adverse metabolic effects of branched-chain amino acids are mediated by isoleucine and valine. Cell Metab, 33(5), 905–922 e906. doi:10.1016/j.cmet.2021.03.025

Yu, D., Tomasiewicz, J. L., Yang, S. E., Miller, B. R., Wakai, M. H., Sherman, D. S., … Lamming, D. W. (2019). Calorie-Restriction-Induced Insulin Sensitivity Is Mediated by Adipose mTORC2 and Not Required for Lifespan Extension. Cell Rep, 29(1), 236–248 e233. doi:10.1016/j.celrep.2019.08.084

Yu, D., Yang, S. E., Miller, B. R., Wisinski, J. A., Sherman, D. S., Brinkman, J. A., … Lamming, D. W. (2018). Short-term methionine deprivation improves metabolic health via sexually dimorphic, mTORC1-independent mechanisms. Faseb J, 32(6), 3471–3482. doi:10.1096/fj.201701211R

Zhang, Y., Xie, Y., Berglund, E. D., Coate, K. C., He, T. T., Katafuchi, T., … Mangelsdorf, D. J. (2012). The starvation hormone, fibroblast growth factor-21, extends lifespan in mice. Elife, 1, e00065. doi:10.7554/eLife.00065

